# Detection of a multi-disease biomarker in Saliva with Graphene Field Effect Transistors

**DOI:** 10.1101/2020.05.22.111047

**Authors:** Narendra Kumar, Mason Gray, Juan C. Ortiz-Marquez, Andrew Weber, Cameron R. Desmond, Avni Argun, Tim van Opijnen, Kenneth S. Burch

## Abstract

Human carbonic anhydrase 1 (CA1) has been suggested as a biomarker for identification of several diseases including cancers, pancreatitis, diabetes, and Sjogren’s syndrome. However, the lack of a rapid, cheap, accurate, and easy-to-use quantification technique has prevented widespread utilization of CA1 for practical clinical applications. To this end, we present a label-free electronic biosensor for detection of CA1 utilizing highly sensitive graphene field effect transistors (G-FETs) as a transducer and specific RNA aptamers as a probe. The binding of CA1 with aptamers resulted in a positive shift in Dirac voltage *V*_*D*_ of the G-FETs, the magnitude of which depended on target concentration. These aptameric G-FET biosensors showed the binding affinity (*K*_*D*_) of ∼ 2.3 ng/ml (70 pM), which is four orders lower than that reported using a gel shift assay. This lower value of *K*_*D*_ enabled us to achieve a detection range (10 pg/ml - 100 ng/ml) which is well in line with the clinically relevant range. These highly sensitive devices allowed us to further prove their clinical relevance by successfully detecting the presence of CA1 in human saliva samples. Utilization of this label-free biosensor could facilitate the early stage identification of various diseases associated with changes in concentration of CAs.

## Introduction

Carbonic anhydrases (CAs) are a family of enzymes that catalyze the reversible reaction of carbon dioxide and water into bicarbonate and protons. ^1^ Several studies have pointed to the importance of the CAs and their inhibitors as an indicator for various diseases.^2–6^ Of particular importance is the human isozyme CA1, which has been identified as a potential marker of various cancers (breast, oral, colorectal, lung, prostate), ankylosing spondylitis, diabetes, Sjogren’s syndrome, and pancreatitis. ^7–12,12–21^ Also encouraging for early detection, the level of CA1 can be determined at point of care (POC) by its presence in easily obtained oral fluid. ^22^ Despite their promise, the current methods for detecting CAs are time consuming, expensive, require extensive clinical processing and expertise in analysis (ELISA, 2D PAGE, and mass spectrometry). ^6^ The lack of high sensitive, rapid, cheap, miniaturized, and user friendly diagnostics has restricted the utilization of CAs for research and clinical purposes.

Graphene Field Effect Transistors (G-FETs) have shown great promise in the detection of DNA, proteins, and cells due to their high sensitivity, scalability, biocompatibility, ease of functionalization via a non-covalent attachment process, and compatibility with various substrates. ^23–31^ To determine the possibility of POC detection of CA, it is crucial to investigate the mechanisms of interaction between G-FETs and CA1, their detection limit, sensitivity, selectivity, and utility in a clinically relevant environment. ^32^ Typically, G-FETs rely on changes in their electrical conductance resulting from the presence of target molecules on the graphene channel. However, one can quantitatively determine the change in the carrier density, and thus target density, by measuring the resulting change in the chemical potential of the G-FET. Since graphene has a minimum in its density of states at the Dirac point, by sweeping an external gate the charge on the sheet is revealed by the voltage (V_*D*_) at which the resistance is maximized. Upon exposure to a target that charges the graphene, a shift in V_*D*_ will occur. As such, the sensitivity of G-FET’s can be limited by their mobility (reduction in the resistance peak height) and Debye screening (reduction in the induced charge on the channel). In the latter case, large probes are particularly detrimental as they keep the target farther from the channel, dramatically reducing the induced charge.

To achieve high sensitivity, we produced clean and reproducible G-FETs by fabricating them in our cleanroom in a glovebox using the same process that produced devices for selective detection of antibiotic resistant bacteria. ^27^ Our facility allows rapid prototyping with minimal reduction in graphene quality from unwanted impurities that subsequently reduce the accuracy and sensitivity. ^33^ Secondly, we chose commercially available RNA aptamers as probes that were screened to be specific for CA1 using the SELEX Method. ^34^ The small size, easy synthesis, and high stability make apatmer probes particularly advantageous for incorporation into G-FET devices. ^35^ Specifically, their smaller size reduces the Debye screening effect, producing high sensitivity (∼ pg/ml) for the detection of proteins utilizing DNA aptamer probes. ^36–38^ Also, the strongly charged backbone of aptamers results in significant changes in the charge carrier density in the graphene channel upon folding or unfolding during target binding. As such, aptamers enable detection in solution with higher ionic strength as desired for real time health care monitoring. ^39^

Here, we present a label-free electronic biosensor for detection of CA1 by utilizing G-FETs functionalized with RNA aptamer probes with high sensitivity, reproducibility, and clinical relevance. Liquid gating was chosen for detection of CA1 to minimize the operating voltage range and keep the aptamers and CA1 in their original structures as well as activities. The shift in V_*D*_ (charge neutrality point) was monitored upon binding of CA1 as it is quantitatively related to the charges induced by binding of targets and not sensitive to the disorder that enhances resistance. Then, two different concentrations of phosphate buffer saline (PBS) were investigated to study their effects on the sensitivity. Furthermore this helped to test whether the change in charge on the graphene resulted from the folding of the aptamers or from the CA1. To determine the sensitivity and detection limit, changes in the V_*D*_ of the G-FET sensors were monitored upon exposure to various concentrations of CA1. The G-FET biosensors showed a lower detection limit of 10 pg/ml within 30 minutes of incubation. The obtained detection limit with this fast, label-free detection method is well below the critical value (∼ 2.6 ng/ml) that has been reported for lung cancer. ^9^ The achieved detection range (10 pg/ml - 100 ng/ml) is comparable to commercially available techniques while eliminating several other steps and without requiring an enzyme tagged antibody, substrates (reagent), and fluori/colori-meters. Negative control tests were also performed to cross-check the selectivity of these aptamers. We found no significant shift of V_*D*_ when the G-FET was exposed to bovine serum albumin (BSA). As a secondary control we tested the effect of direct exposure of the G-FETs to CA1 without aptamers, finding no net shift in the V_*D*_. To further demonstrate the clinical relevance of G-FETs and their utilization for future POC diagnostics, human saliva was chosen as a clinical body fluid. Salivary bio-markers for disease detection are of increasing interest because the collection of saliva samples is non-invasive, painless, and achievable with minimal training. ^40–42^ The G-FET biosensors enabled the successful detection of unknown concentrations of CA1 present in obtained human saliva. Furthermore, similar to our observations in PBS, the increase in output signal (V_*D*_) was observed when the saliva was spiked with higher concentrations of CA1, confirming the detection capability of the presented G-FET biosensors in clinically relevant samples.

## Materials and Methods

### Device Fabrication

The G-FET devices were fabricated on commercially available CVD-grown monolayer graphene on *SiO*_2_*/Si* substrates obtained from Graphenea. The graphene was baked at 300 *°*C for 9h in vacuum to remove photoresist or other impurities in addition to improving its adhesion with the substrate. To protect the clean graphene from exposure with resist and chemicals during fabrication, 3 nm thick AlO_*x*_ was deposited by evaporating aluminum in an oxygen pressure of 7.5 × 10^−5^ mbar. After deposition, the substrates were transferred to an Argon atmosphere in a glove box for fabrication and baked at 175*°*C for 5 minutes for better adhesion of the AlOx layer. The electrode patterning was done using bilayer photoresist (LOR1A/S1805) and a maskless lithography system (Heidelberg Instruments) followed by Au/Cr (45 nm/5 nm) deposition and lift off using remover PG. The graphene was patterned to define the active region using photolithography and oxygen plasma etching. The remaining AlOx layer was etched away by submerging the chips in fresh developer for 60 s. In order to protect the electrodes and edges of the graphene for liquid gating, 20 nm AlOx was deposited and baked at 175 *°*C for 10 minutes. Then to open up the sensing window of desired size (6*µm* × 20*µm*) and contact pads,the patterning was obtained by lithography using S1805 and then Transetch-N was utilized for AlOx removal. Etching was performed on a 70 *°*C hot plate for an optimized time of four minutes followed by resist removal with remover PG, rinsing with isopropyl alcohol (IPA), DI water, and dried with Argon. A typical device structure and a microscopic image are shown in Figure 1a and 1b, respectively. A PDMS well with 40*µl* volume was mounted on the chips to hold solutions in place.

**Figure 1:**
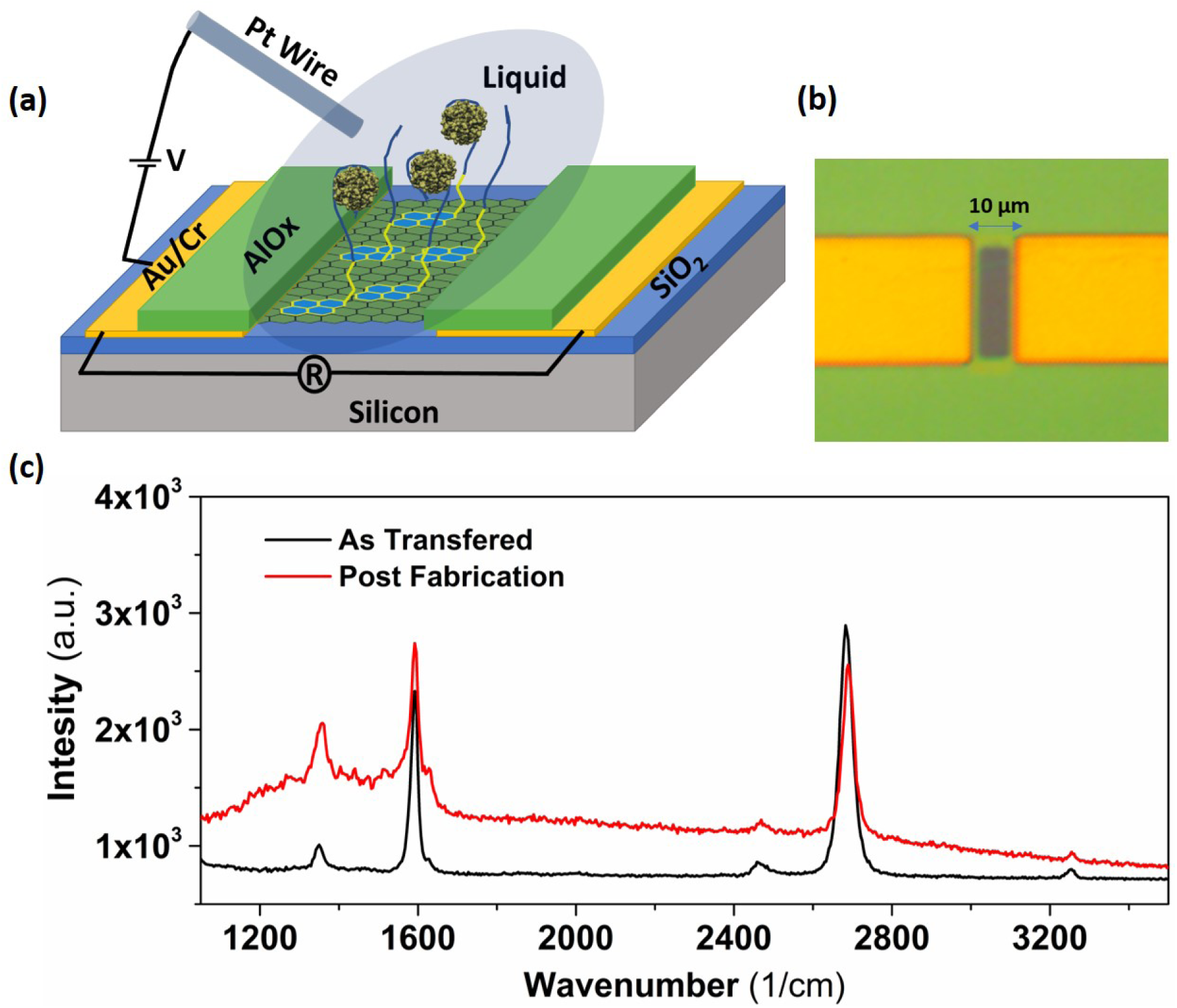
(a) Schematic representation of a G-FET device utilized for biosensing consists of graphene over SiO_2_/Si substrate, Au/Cr (45/5 nm) source/drain electrodes, AlOx layer (20 nm) for passivation to define the sensing area, Aptamers attached to graphene through pyrene linker, CA1 binding resulting in folding down of aptamer probes, and a Pt reference electrode for liquid gating. (b) Microscopic image of a fabricated G-FET, having a graphene sensing window (6 *µ*m x 20 *µ*m) made from patterned AlOx also acting as a passivation layer for Au/Cr electrodes. (c) Raman spectra of graphene before and after fabrication. The G band (1591 *cm*_1_) is at the same position suggesting fabrication does not affect the doping level of graphene, while a slight increase in background and enhanced D peak suggests some additional, charge neutral disorder.

### Biological and Chemical Materials

1-pyrenebutyric acid N-hydroxysuccinimide ester (PBASE) linker and Dimethylformamide (DMF) were obtained from Sigma Aldrich. Saliva samples were purchased from ProMedDx, LLC (Norton, MA). Samples were collected via passive drool technique from a minimum of 5 apparently healthy subjects. Collected samples were aliquoted into 0.25 mL volume and Immediately frozen at -80*°*C. No preservatives or additives were used in the sample. The saliva spiked with CA1 was collected from the same subject and homogenized before use. 5 end amine modified RNA aptamers with sequence CAGGCGCAGUAUCACUUUGUGAUCAUUAGUGGGUUCCGUG were obtained from Integrated DNA Technologies.

## Results and discussion

### G-FET Characterization

To ensure the uniformity and high quality of the fabrication, each device was characterized by Raman and electrical measurements. Figure 1c shows the Raman characterization of the graphene before and after fabrication. As received graphene showed G (1591 *cm*^−1^) and 2D (2682 *cm*^−1^) bands with a 2D/G ratio of 1.4, confirming single layer graphene. ^43^ As shown in Figure 1c, no change was observed in the position of G band after fabrication, though a slight increase in the intensity of the D peak (1358 *cm*^−1^) along with some background absorption was observed. This suggests the fabrication did not significantly alter the doping level and causes minimal disorder in the graphene. ^44^ The reproducibility and mobility of the fabricated G-FETs was determined by measuring the two point (between source and Drain) resistance while sweeping the liquid gate voltage. The devices fabricated in five different batches showed an average *V*_*D*_ of 0.7 ± 0.1*V* measured at 0.01x PBS using Pt wire as reference electrode for gating. The non-zero value of *V*_*D*_ is attributed to the surface potential associated with Pt wire and not due to doping from the fabrication process. ^31,45^ The obtained average field effect mobilities for electrons and holes were *µ*_*n*_ = 1230 *cm*^2^*/V.s* and *µ*_*h*_ = 1040 *cm*^2^*/V.s* respectively, demonstrating the high quality of our process and CVD graphene. ^46,47^ However, we have seen variation in the values of the maximum resistance measured with devices made in different chips/batches which could be either due to the disorder in the graphene or contact resistance. Therefore for all the sensing experiments, we ignored the value of resistance and only focused on the shift in *V*_*D*_ which is expected to only be affected by change in doping level.

### G-FET functionalization

In order to achieve the selective detection of CA1 enzyme, the G-FETs need to be functionalized with respective aptamers. Further, to prevent false positives that could occur due to nonspecific adsorption of CA1, it is crucial to obtain uniform functionalization of aptamers on the graphene surface. As such, the attachment process was optimized using atomic force microscopy (AFM) to determine attachment and coverage of the aptamers. ^27^ Patterned graphene chips were incubated for 30 minutes with high concentration (10 mM) PBASE linker dissolved in DMF.^25,31^ Next, the G-FET was rinsed with DMF to remove adsorbed linker molecules followed by rinsing with IPA, DI to make the surface free from solvents. The pyrene group in PBASE linker stacks over graphene surface through *π* −*π* interaction while the N-hydroxysuccinimide (NHS) ester reacts with amine terminated at 5’ end of aptamers. ^23^ Chips functionalized with linker were incubated for 30 minutes with PBS solution containing aptamers with a concentration of 1*µM*. ^25,31^ Finally, the chips were rinsed with PBS to remove excess aptamers, followed by DI to clean the salts from the graphene surface and finally dried with Argon for AFM characterization. Figure 2 confirms the attachment and complete coverage of the aptamers as the thickness of the functionalized graphene is uniformly ∼ 4 nm with respect to SiO_2_ surface. The change in thickness of the functionalized graphene was measured at several different locations while single step is shown as a representative plot. The relative increase in height from bare graphene is ∼ 2.5 − 3 nm and is consistent with earlier reports. ^31,36,46^ Some dots were also seen on the SiO_2_ surface probably due to the nonspecific adsorption of linker or aptamers while the untreated SiO_2_ surface was clean (see Figure 2a). The attachment of aptamers was further confirmed by Raman characterization. Several peaks (1235*cm*^−1^, 1350*cm*^−1^, 1387*cm*^−1^,1527*cm*^−1^, 1624 *cm*^−1^) emerged upon functionalization of graphene with aptamers (Figure 2 b). The observed peaks at 1235*cm*^−1^, 1387*cm*^−1^, 1624 *cm*^−1^ are attributed to the PBASE linker while other two at 1350*cm*^−1^, 1527*cm*^−1^ are likely due to the aptamers attachment. ^25,48^ The decrease in the ratio of 2D/G is another confirmation of the surface modification of graphene. We note these measurements were taken after the substrates had been rinsed and dried, further confirming the attachment of the linker and aptamer to graphene. After confirming the aptamers’ functionalization on bare graphene, we moved to ensure the attachment process also works for fully fabricated devices. To do that, the G-FETs were incubated with PBASE linker and aptamers using the same protocol used for bare graphene. The change in *V*_*D*_ was then monitored after functionalization. As expected from the negatively charged backbone of aptamers, ^39^ a significant positive shift in *V*_*D*_ was observed (see Figure 3), confirming the attachment of aptamers on the G-FETs.

**Figure 2:**
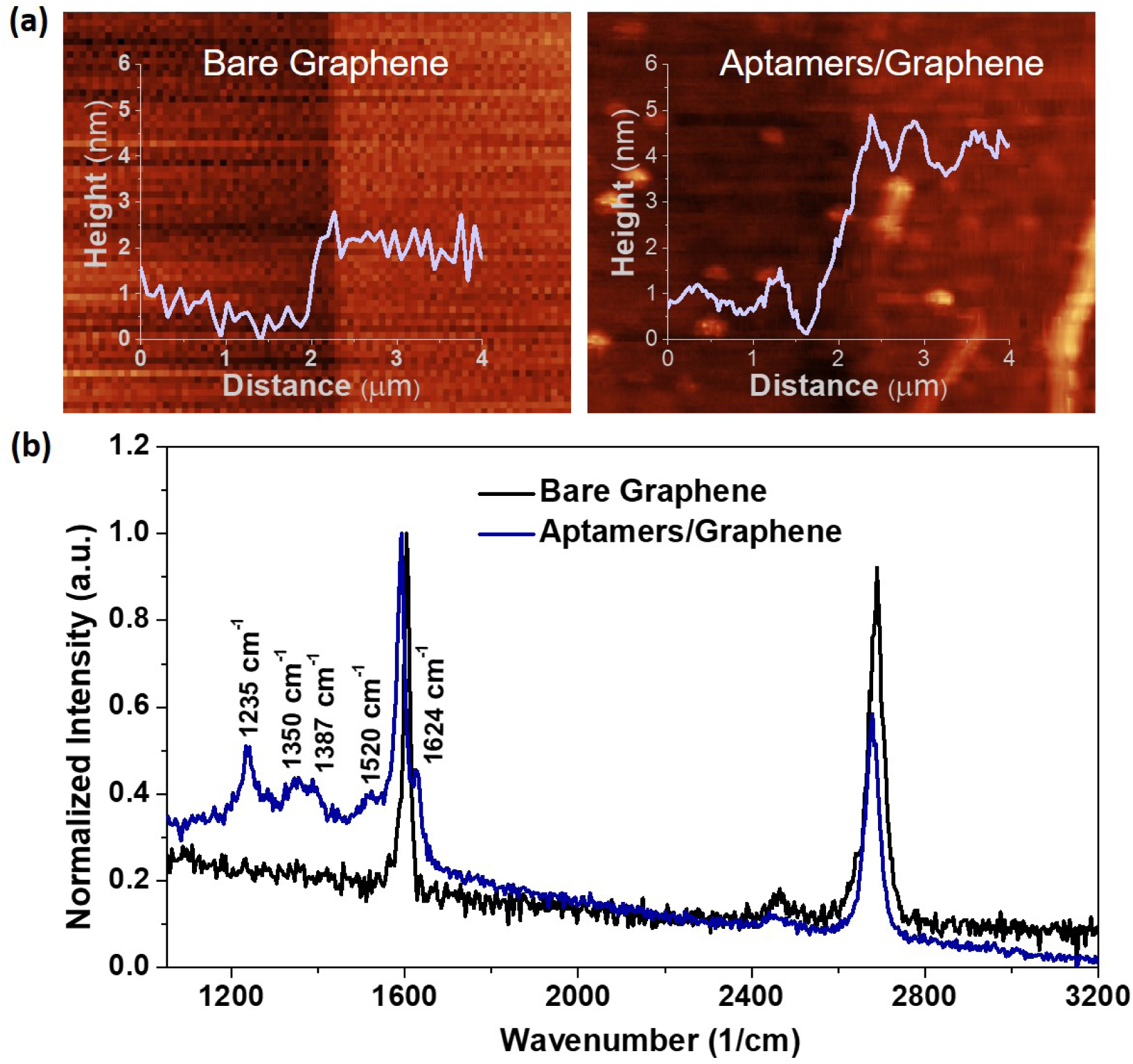
(a) AFM image of bare graphene (left) and after attachment with aptamer (right). A change in step height was measured when AFM tip travels from *SiO*_2_ surface to Graphene/*SiO*_2_. The ∼ 2.5 nm increase in thickness of the functionalized graphene with aptamers confirms the attachment. Furthermore, the uniformity of the height increase indicates full coverage. (b) Raman spectrum of graphene before and after functionalization. Emergance of several peaks (1240*cm*^−1^, 1350*cm*^−1^, 1389*cm*^−1^,1527*cm*^−1^, 1624 *cm*^−1^) further confirming the modifications of graphene surface with linker and aptamer.

**Figure 3:**
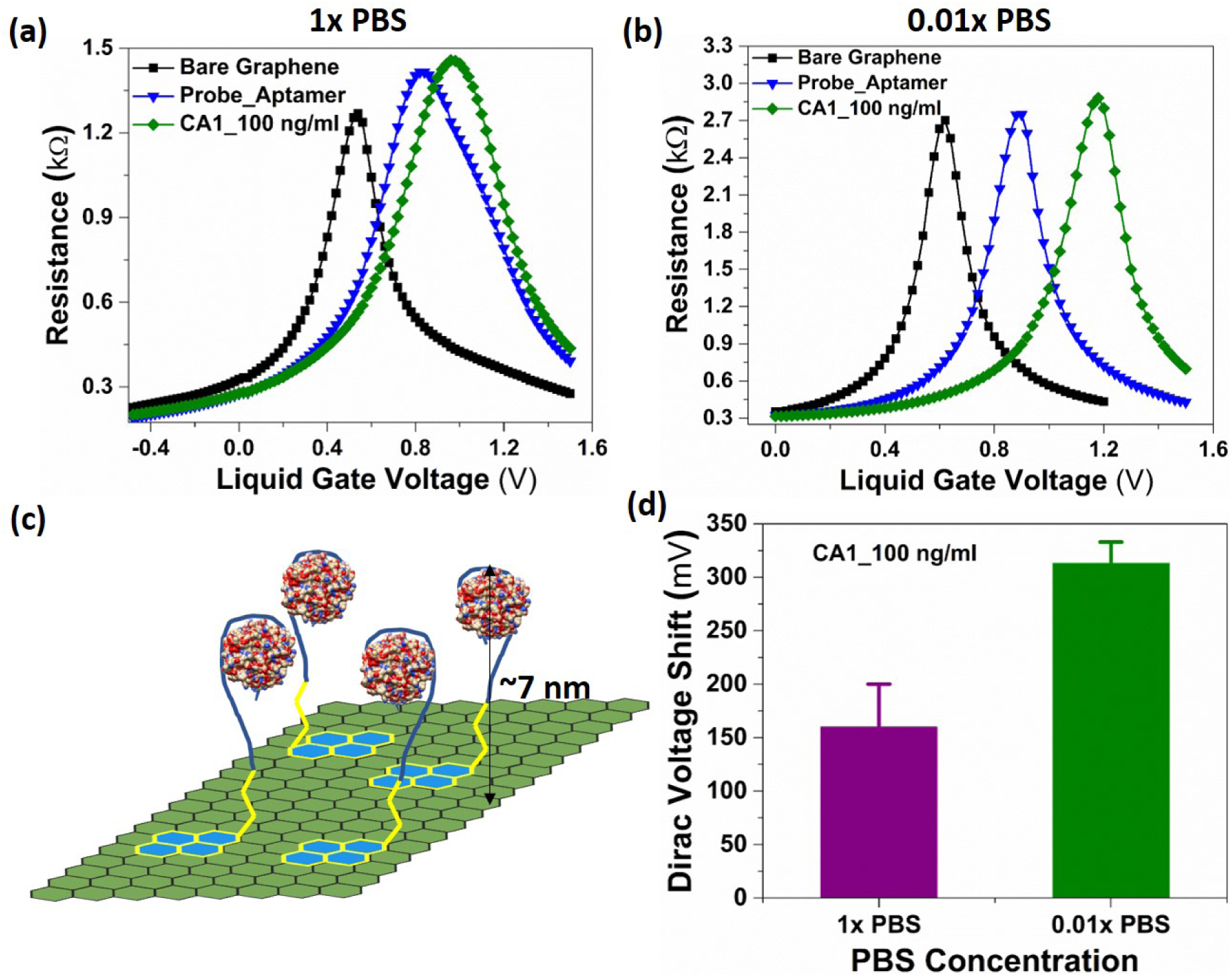
Effect of aptamer and CA1 binding on G-FET characteristics (a) Incubation of CA1 (100 ng/ml) and measurement in 1x PBS, a shift of ∼ 160*mV* was observed, indicating folding down of aptamers upon binding with CA1 (b) Incubation of CA1 in 1x PBS and measurement in 0.01x PBS, resulting in higher shift of ∼ 313*mV* resulting from a combination of charges on CA1 and folding down of aptamers. (C) Schematic of CA1 binding on graphene surface, illustrating approximate distance of CA1 from G-FET surface. (d) Comparison of signal measured in 1x PBS and 0.01x PBS, revealing a 2-fold increase in output signal for the same CA1 concentration.

### Optimizing the Buffer

After confirming the aptamers’ functionalization on G-FETs, we turned to investigate and maximize the sensitivity of the devices to CA1. Given the size of the aptamers (∼ 2.5 nm see Figure 2) and the CA1 (∼ 4.25 nm), ^49,50^ it is crucial to carefully consider the Debye screening length of the employed buffer. Specifically, if the Debye length is too small it is likely to dramatically reduce the sensitivity of the graphene to the change in charge density induced upon binding of CA1 with the aptamers. Therefore, the measurements were performed with two different buffer strengths, 1x PBS and 0.01x PBS whose Debye lengths are 0.75 nm and 7.3 nm, respectively. Since the diluted PBS has a Debye length longer then the aptamers and target, we expect significantly larger signals than with full strength PBS which almost completely screens the charge. In spite of expected lower signals, the physiological relevance of full strength buffers make them crucial for real-time healthcare monitoring. Therefore, we first performed the measurement in 1x PBS using CA1 concentration of 100 ng/ml. As shown in Figure 3a, a positive shift in *V*_*D*_ was observed upon functionalization of graphene with aptamers which is attributed to their negatively charged backbone. Upon exposure to CA1, an additional shift was also observed. Considering the 1x PBS Debye length of 0.75 nm and size (∼ 2.5 nm) of the aptamers, no shift in *V*_*D*_ was expected. However a shift of 160 mV in *V*_*D*_ suggesting some conformational changes in the aptamers, i.e. folding which brings the aptamers closer to the graphene surface upon binding with CA1. This observation supports the hypothesis reported for the detection of dopamine and glucose using an aptamer based *In*_2_*O*_3_ FET sensor. ^39^ Thus the obtained detection of CA1 in 1x PBS utilizing aptamer probes could enable G-FET devices to detect analytes in physiological solutions and their subsequent utilization for real-time monitoring. Moreover, if the shift is caused by the folding of aptamers and independent of charges on the target, it can also be utilized for neutral analytes.

Though the G-FETs can detect the CA1 in 1x PBS, the small shift relative to that caused by the aptamers could limit the detection of target concentrations too low to be clinically relevant. Therefore, we moved to maximize the signal by measuring at low buffer strength while incubating (binding) in 1X PBS. Knowing that CA1 possesses a negatively charged surface at pH 7^51^ because of its isoelectric point of 6.67, ^52^ there is scope to further enhance the obtained shift in *V*_*D*_ by increasing the Debye length to 7.3 nm using 0.01x PBS. To achieve this, the measurements were conducted at diluted 0.01x PBS as shown in Figure 3b. As expected, the obtained shift in *V*_*D*_ upon binding of CA1 was approximately twice as large as that observed at 1x PBS (see Figure 3d). The two-fold more positive shift in *V*_*D*_ upon binding with CA1 is attributed to the combination of charges induced by CA1 and folding down of aptamers closer to the sensor surface. Since the estimated size of aptamer-CA1 assembly is ∼ 7*nm* (Figure 3c), the screening has little effect in 0.01x PBS possessing a Debye length of 7.3 nm, resulting in a higher output signal of ∼ 313*mV* (Figure 3d).

### Specificity of Detection

Having optimized the device functionalization and measurement conditions, we turned to utilize the G-FETs for specific and quantitative detection of CA1. Based on the observations of higher output signal, the 0.01x PBS was used during the measurement while the incubation of aptamers as well as CA1 was carried out in physiologically relevant 1x PBS. In order to test the specificity of these devices, we first measured the effect of CA1 without attaching aptamers (i.e. on unfunctionalized G-FETs). As shown in Figure 4a, very little shift (∼ 40 mV) was observed when the G-FET was exposed to CA1 without aptamers, similar to the shift seen when only PBS was applied. The same device showed significant shift after first attaching the aptamers and then reexposing to CA1 (Figure 4a). This confirmed that CA1 can not attach to the graphene surface without functionalization of aptamers. To further validate the selectivity of aptamers for CA1, the devices functionalized with aptamers were incubated with 100 ng/ml BSA and an average shift of ∼ 13*mV* was observed as shown in Figure 4b (red circle), a value within the standard deviation of simply applying 1x PBS and below that obtained with four orders of magnitude lower concentration of CA1.

**Figure 4:**
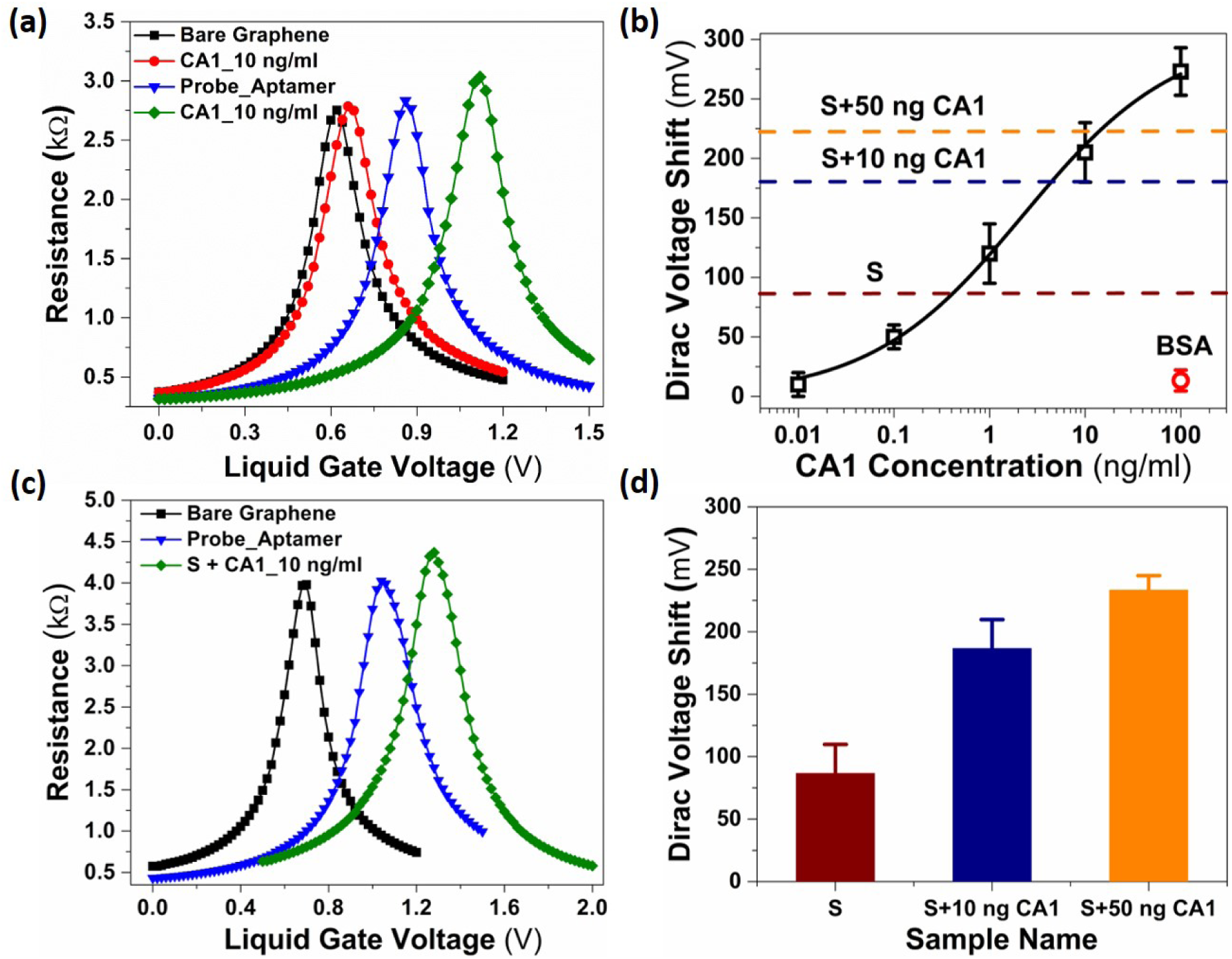
Determination of G-FETs’ sensitivity and specificity of CA1 in PBS (a) Effect of CA1 detection on the G-FET with and without aptamers. A minimal shift (∼ 40 mV) in *V*_*D*_ is observed at CA1-10 ng/ml measured on the bare G-FET, while ∼ 220 mV was measured after the use of aptamers. This confirms CA1 is only binding with aptamers and not adsorbing specifically on the graphene surface. (b) Average shift in *V*_*D*_ tested at different concentrations of CA1, using 0.01x PBS for gating. Additionally the minimal shift with BSA at the top concentration used for CA1, confirms the specificity of the aptamers. The solid line is a fit to Hill’s equation 1, revealing high binding affinity (K_*D*_) of 2.27 ng/ml with a detection range of 10*pg/ml < ρ <* 100*ng/ml*. Dashed lines indicate V_*D*_ measured for as received (S), and spiked human saliva diluted in ratio 1:10 in 1x PBS. (c) G-FET characteristics showing shift in *V*_*D*_ measured with saliva + CA1-10 ng/ml (d) Average shift in *V*_*D*_ with as received saliva and that spiked with two different concentrations of CA1.

After demonstrating the selectivity of the devices, we moved to determining the sensitivity and lower detection limit of G-FETs functionalized with aptamer probes. Specifically, we measured the response of the G-FETs to concentrations of CA1 ranging from 100 ng/ml to 10pg/ml. This was done with a minimum of three devices for each concentration and different devices were used to test different concentrations. Each device was calibrated by first determining the Dirac Voltage in diluted 0.01x PBS (*V*_*D*_(0)). The resulting change in Dirac voltage (Δ*V*_*D*_(*ρ*) = *V*_*D*_(*ρ*) − *V*_*D*_(0)) of the G-FET biosensor due to the exposure to CA1 of a given concentration (*ρ*) is shown in Figure 4b. The expected increase in voltage shift was observed with increasing concentration of CA1 ranging between 10*pg/ml < ρ <* 100*ng/ml*. Considering the molecular weight of CA1 (33 kDa), ^49^ the measured concentration range is equivalent to 330*f M < ρ <* 3*nM*. The achieved lower detection limit of 10 pg/ml is well below the reported critical value of ∼ 2.6 ng/ml indicative of non-small cell lung cancer. ^9^

In order to analyse the performance of the G-FET biosensor, the sensitivity was calculated by linearly fitting the plot of Δ*V*_*D*_ versus CA1 concentration; the obtained slope of the plot reveals a sensitivity of 65.4 mV/dec with a linearity (*R*^2^)of 0.97. Furthermore, to better understand the binding kinetics of the aptamer probes with CA1, a fit of the data to a calibration curve was obtained by using Hill’s equation which is well known to describe the ligand-receptor interaction:^53^

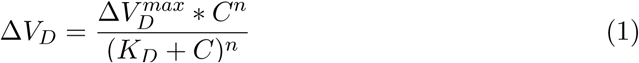

where ΔV_*D*_, is the measured Dirac voltage shift at different concentration of CA1, 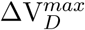 is the Dirac voltage shift when all the binding sites are saturated, C is the concentration of CA1, K_*D*_ is the dissociation constant, and n is the Hill’s coefficient. As shown in Figure 4b, the resulting fit provides an excellent description of the concentration dependence. Interestingly, we find a K_*D*_ of 2.27 ng/ml (∼ 70 pM), which is four orders lower than (386 nM) obtained using a gel shift binding assay with the same aptamer probes. ^34^ Moreover, the obtained value is consistent with that reported in the literature for different aptamers and bio-analytes using G-FET devices ^37,38^ while one to three orders lower than those obtained with any other optical or mass based technique. ^35^ We find 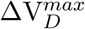 of 305.4 mV consistent with our experience that devices with CA1 concentration above 100 ng/mL did not produce additional shifts. The value of n (0.55) is in agreement with an aptamer designed with single binding sites. ^54^ A value lower than 1 is attributed to the negative cooperativity of receptor-ligand binding. ^55^ The higher binding affinity (2.27 ng/ml) of our sensor can be explained in two ways. As suggested by Ha et al., ^56^ the negative cooperativity is explained by one receptor binding with two ligands, producing higher binding affinity. This suggests that the CA1 enzyme can simultaneously bind with two of the aptamers used in our study. Another possible explanation could be on the basis of heterogeneous adsorption isotherm in which the distributed binding energies of aptamer-CA1 interactions are considered. ^24^

### Detection in Saliva

Having confirmed that the G-FET devices functionalized with the designed RNA aptamer are highly sensitive and selective to CA1 enzyme, we turned to real clinical samples to meet the requirement of practical applications. ^32^ Considering the feasibility of collection and processing, a clinical human saliva sample was chosen. Furthermore, the presence of the CA1 in saliva samples is expected to indicate the presence of Sjogren’s syndrome. ^18–20,42^ To perform the measurements, as received human saliva sample was diluted to a 1:10 ratio in 1x PBS to reduce its viscosity, enabling the solution to easily reach to the sensor surface. Next, the saliva was spiked with CA1 using two different concentrations of 10 ng/ml and 50 ng/ml, those expected to be higher than that already present in the received samples. The G-FET devices functionalized with RNA aptamers were tested with as received and CA1 spiked saliva samples and the respective obtained resistance versus voltage characteristics are shown in Figure 4c. As shown in Figure 4d, as received human saliva sample also produced a voltage shift of 86.6 ± 23 mV, confirming the expected presence of some CA1 in as received healthy human samples. The obtained shift is equivalent to a CA1 concentration of 0.4 ng/ml determined from our 0.01x PBS measurements (Figure 4b), which means the sample contained the CA1 concentration of 4 ng/ml as it was diluted 10 times for the measurement. To confirm this shift is does not result from molecules in saliva, other than CA1, the G-FET biosensor was tested with saliva spiked with higher concentrations (10 ng/ml) of CA1. As expected, a further shift in *V*_*D*_ was observed upon incubation with spiked saliva with externally added CA1, confirming the sensor’s sensitivity to CA1. However, the obtained shift with CA1 spiked saliva (Figure 4d) is slightly lower than that observed in PBS (Figure 4b) which can be attributed to the interference of other ions/species present in saliva samples. Furthermore to confirm the concentration dependent sensitivity, we tested with saliva spiked to a higher concentration of 50 ng/ml, resulting in an additional shift in *V*_*D*_ as expected from increased concentration. Also due to the high viscosity of saliva samples, we found that higher incubation time is required. However this only required an increase from 30 minutes to 1 hour for the transportation of the CA1 enzyme to the sensors surface.

## Conclusion

In Summary, we demonstrated G-FET biosensor utilizing RNA aptamer probes showing high sensitivity (65.4 mV/dec) and specificity to a targeted multi-disease biomarker, the CA1 enzyme. The achieved lower detection limit 10 pg/ml with a detection range between 10 pg/ml - 100 ng/ml utilizing this label-free electronic detection method is comparable with the existing ELISA-based commercially available techniques. Furthermore, devices exposed to human saliva samples clearly identified elevated levels of CA1, as required for clinical relevance. Nonetheless future studies are required to ensure minimal sensitivity to other enzymes and bioactive molecules. The G-FET biosensor we developed can detect the CA1 in 30 minutes in PBS and 60 minutes in saliva samples, requiring only 40 *µl* of sample. This study opens the door to explore this method for detection of different carbonic anhydrases and other similar biomarkers by designing specific RNA aptamers. Fast, sensitive, and label-free detection utilizing this platform could speed up the identification of different diseases related with CAs and their early stage detection. Furthermore, the successful demonstration of G-FET biosensors for detection of CA1 in saliva paves the way to identify various diseases using oral fluid.

## Acknowledgement

and K.S.B. acknowledge the primary support of the US Department of Energy (DOE), Office of Science, Office of Basic Energy Sciences under award no. DE-SC0018675. J.O-M and T.v.O were supported by the National Institutes of Health, through grants R01-AI110724 and U01-AI124302. The work of M.G. was enabled by support from the National Science Foundation, through grant DMR-1709987. A.W. and A.A. acknowledge the support from National Institutes of Health/ National Institute of Dental and Craniofacial Research, National Institute on Drug Abuse through grants R43-DE025153 and R43-DA051105.

## Graphical TOC Entry

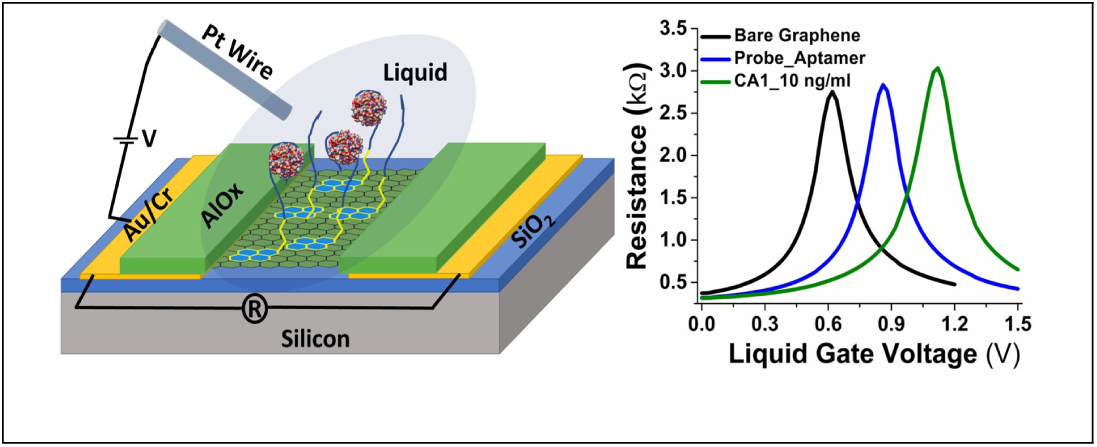

